# A new Bayesian methodology for nonlinear model calibration in Computational Systems Biology

**DOI:** 10.1101/633180

**Authors:** Fortunato Bianconi, Lorenzo Tomassoni, Chiara Antonini, Paolo Valigi

## Abstract

Computational modeling is a common tool to quantitatively describe biological processes. However, most model parameters are usually unknown because they cannot be directly measured. Therefore, a key issue in Systems Biology is model calibration, i.e. estimate parameters from experimental data. Existing methodologies for parameter estimation are divided in two classes: frequentist and Bayesian methods. The first ones optimize a cost function while the second ones estimate the parameter posterior distribution through different sampling techniques. Here, we present an innovative Bayesian method, called Conditional Robust Calibration (CRC), for nonlinear model calibration and robustness analysis using omics data. CRC is an iterative algorithm based on the sampling of a proposal distribution and on the definition of multiple objective functions, one for each observable. CRC estimates the probability density function (pdf) of parameters conditioned to the experimental measures and it performs a robustness analysis, quantifying how much each parameter influences the observables behavior. We apply CRC to three Ordinary Differential Equations (ODE) models to test its performances compared to the other state of the art approaches, namely Profile Likelihood (PL), Approximate Bayesian Computation Sequential Monte Carlo (ABC-SMC) and Delayed Rejection Adaptive Metropolis (DRAM). Compared with these methods, CRC finds a robust solution with a reduced computational cost. CRC is developed as a set of Matlab functions (version R2018), whose fundamental source code is freely available at https://github.com/fortunatobianconi/CRC.

## Introduction

In recent years *omics* technologies have tremendously advanced allowing the identification and quantification of molecules at the DNA, RNA and protein level [1, 2]. These high-throughput experiments produce huge amounts of data which need to be managed and analyzed in order to extract useful information [3]. In this context, mathematical models play an important role since they process these data and simulate complex biological phenomena. The main purpose of mathematical modeling is to study cellular and extracellular biological processes from a quantitative point of view and highlight the dynamics of cellular components interactions [4]. Moreover, models represent an excellent tool to predict the value of biological parameters that may not be directly accessible through biological experiments because they would be time consuming, expensive or not feasible [5]. One of the most common modeling techniques consists in representing a biological event, such as a signaling pathway, through a system of Ordinary Differential Equations (ODEs), which describes the dynamic behavior of state variables, i.e. the variation of species concentration in the system as a function of time [6]. Currently, the most used kinetic laws in ODE models can be divided into three types: the law of mass action, the Michaelis-Menten kinetic and the Hill function [7, 8]. However, these equations contain unknown parameters which have to be estimated in order to properly simulate the model and represent the problem under study. Typically, the calibration process of a model consists in the inference of parameters in order to make output variables as close as possible to the experimental dataset [9]. Hence, a calibrated model can be used to predict the time evolution of substances for which enough information or measures are not available. The most common methodologies for parameter estimation can be divided in two classes: the frequentist and the Bayesian approach [5, 10]. Frequentist methods aim at maximizing the likelihood function *f* (**y**|**p**), which is the probability density of observing the dataset **y** given parameter values **p** [11]. Under the hypothesis of independent additive Gaussian noise with constant and known variance for each measurement, the Maximum Likelihood Estimation (MLE) problem is equivalent to the minimization of an objective function, which compares simulated and experimental data [12, 13]. Common objective functions are the sum of squared residuals or the negative log-likelihood [14]. When the variances of measurement noise are not known, they are included in the likelihood as additional terms to estimate [13]. Different optimization algorithms are then employed to estimate the best parameter values. They implement global and/or local techniques and return in output the best fit between simulated and real data.

Since these optimization methods return only one solution for the parameter vector, i.e. the best fit, then parameter estimation is usually combined with identifiability analysis, in order to assess how much uncertainty there is in the parameter estimate [15]. Identifiability analysis is typically performed through the computation of confidence intervals for all estimated parameters. A confidence interval is the range where the true parameter value is located with a certain frequency. In this context, Profile Likelihood (PL) is a widely used data-based algorithm for structural and practical identifiability analysis [16].

On the other hand, the Bayesian approach considers parameters as random variables, whose joint posterior distribution *f* (**p**|**y**) is estimated through the Bayes theorem: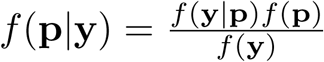, where f(**y**) is the marginal density for **y** and f(**p**) is the prior probability density of parameters [17]. Through f(**p**) it is possible to include *a priori* beliefs about parameter values [18]. The joint posterior density automatically provides an indication of the uncertainty of the parameter inference and gives major insights about the robustness of the solution [19]. Since computing the posterior distribution analytically is usually not feasible, sampling based techniques are used to estimate it [12, 17]. Two classes of sampling methods widely used are the Markov chain Monte Carlo (MCMC) and the Approximate Bayesian Computation Sequential Monte Carlo (ABC-SMC) [12]. MCMC algorithms approximate the posterior distribution with a Markov chain, whose states are samples from the parameter space. Their major advantage is the ability to infer the posterior distribution which is known only up to a normalizing constant [20]. The ABC-SMC algorithms evaluate an approximation of the posterior distribution through a series of intermediate distributions, obtained by iteratively perturbing the parameter space. Each iteration selects only those parameters that give rise to a distance function under a predefined threshold [21].

In this paper, our main purpose is to introduce a new version of the standard ABC-SMC approach, called Conditional Robust Calibration (CRC), for parameter estimation of mathematical models. As in all ABC-SMC methods, CRC is an iterative procedure based on the parameter space sampling and on the estimation of the probability density function (pdf) for each model parameter. However, it presents different aspects that differentiate it from the other existent methodologies. The distinctive features of CRC are: (i) a major control of the computational costs of the procedure, (ii) the definition of multiple objective functions, one for each output variable, (iii) the conditional robustness analysis (CRA) [22], in order to determine the influence of each model parameter on the observables.

We validate this new methodology on different ODE models of increasing complexity. Here we present the results of CRC applied to three models, two with *in silico* noisy data and one with experimental proteomic data. We also calibrate all the models using the PL approach, through the software Data2Dynamics (D2D) [23], the standard ABC-SMC, through the ABC-SysBio software [19] and an MCMC algorithm called Delayed Rejection Adaptive Metropolis (DRAM) through the MCMC toolbox [24, 25], with the purpose of having a reliable and complete comparison with the state of the art of this field [24]. Our results show that CRC is successful in all examples, with several advantages when compared to the other methodologies.

## Materials and methods

### ODE model and experimental dataset

Consider a deterministic ordinary nonlinear dynamical system:

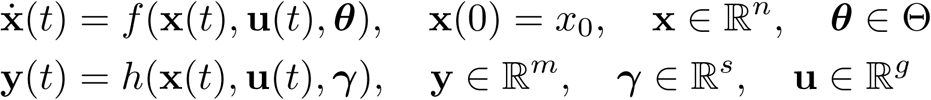

where **x** is the state space vector, **u** denotes the external input vector, and **y** denotes the output responses of the system, i.e. the observables, which are usually derived from experimentally observed data. The vector ***θ*** denotes the dynamical system parameters, taking values in the parameter space **Θ**, a subset of the positive orthant 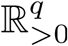. The vector function *f* is indeed defined over the following sets: *f* (·): ℝ^*n*^ × ℝ^*g*^× ℝ^*q*^ → ℝ^*n*^. The observation function *h*(·): ℝ^*n*^ × ℝ^*g*^ × ℝ^*s*^ → ℝ^*m*^ maps the state variables to the vector of observable quantities **y**. Usually, not all states of the system can be directly measured, so that it is common to have *m < n*. Vector *γ* contains scaling and offset parameters when measurements of the observables are performed. Setting **p** = {*θ, γ,* **x**(0)}, **p** ∈ ℝ^*q*+*s*+*n*^, the model is completely determined. We assume that the parameter vector **p** is constant over time.

### General theory

The parameter vector of a mathematical model can be considered as a random variable **P** on a measurable space (ℙ, *A*) and a given vector **p** in the parameter space as one of its realizations. Let *f***_P_**(**p**) denote the prior distribution. Our goal is to approximate the target posterior distribution, *f*_**P**|**y**_(**p**)∝*f* (**y|p**)*f***_P_**(**p**), where *f* (**y|p**) is the likelihood density that describes the model. To this purpose, we develop CRC, a variant of the ABC-SMC iterative algorithm. As it is well-established in this class of techniques, at the beginning of each iteration *z*, CRC samples parameters from a proposal distribution *q*^*z*^(**p**). However, differently to standard ABC approaches, we generate a fixed number *N*_*S*_ of samples from *q*^*z*^(**p**) along the different iterations. Then the fitting between observed and simulated data is measured through the computation of a distance function. CRC defines, at each iteration, multiple distance functions *d*_*i,p*_, each one associated with a single component of the output function, without the employment of any summary statistic. Since parameter vector **P** is a random variable, *d*_*i,p*_ can be considered a realization of the random variable *D*_*i*_ that describes the distribution of the distance function, corresponding to the *i-th* output. Accordingly, at each iteration we define a set of thresholds 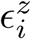 that specify the maximum accepted level of agreement between each observable and the corresponding simulated data. At each iteration we obtain different parameter sets 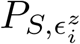. Each set contains only those parameters that yield the values of a specific distance function under the corresponding threshold. Then, all these sets are intersected in order to obtain a single parameter set, 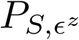, that ensures the compliance with all the thresholds.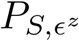 contains samples of the approximate posterior distribution 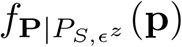 (**p**). As for other ABC methods, if at the end of the *z-th* iteration a predefined stopping criterion is not satisfied, another iteration of CRC is performed, sampling from a new proposal distribution. Since the region of interest of the approximate posterior distribution is the one with highest probability, the proposal for the next iteration, *q*^*z*+1^(**p**), is centered on the mode of 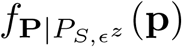. In order to increase the frequency of *N*_*S*_ samples in this region, the boundaries of *q*^*z*+1^(**p**) are tighter than those of *q*^*z*^(**p**). The algorithm terminates when the thresholds are sufficiently small. The output of CRC is 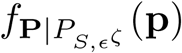 where *ζ* is the number of the last iteration and its mode 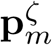 is the vector that reproduces the observed data with the highest probability.

### CRC algorithm

Fig 1 sums up the main steps of the CRC algorithm. **1.Sampling the parameter space.** Generate a predefined number of samples *N*_*S*_ from the proposal distribution *q*^*z*^(**p**). If *z* = 1 the proposal distribution is the prior *f***_P_**(**p**).

**Fig 1.**
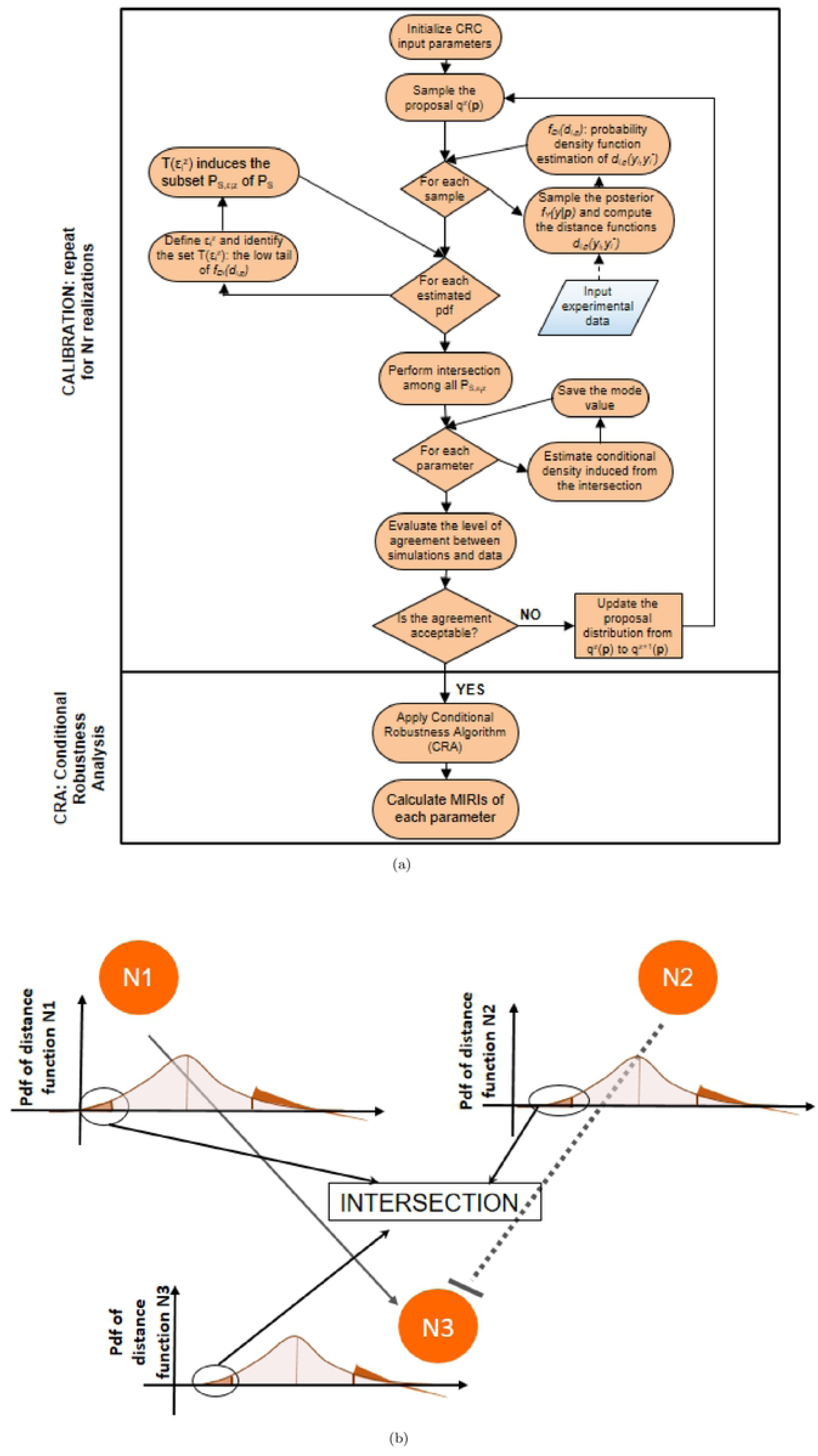
CRC algorithm. The flowchart of CRC is divided in two main phases: model calibration and robustness analysis.

**2. Sampling the posterior distribution.** Let *P*_*S*_ be the set of parameter samples generated in the previous step. For each **p** ∈ *P*_*S*_, sample *y*_*i*_, ∀*i* = 1, …, *m* from the posterior distribution *f***_Y_**(**y**|**p**).

**3. Computation of the distance functions and pdf estimation.** For each **p** ∈ *P*_*S*_, as many distance functions as observables are computed, i.e.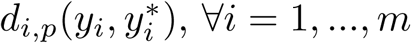, where 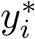 is the *i-th* variable of the experimental dataset. Then the associated densities are estimated using a kernel density approach. Let denote with 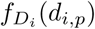 the pdf of *D*_*i*_. **4.Parameter sets identification.** For each distance function we define a threshold 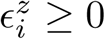 which is the quartile of level *α* of 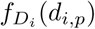 and we obtain the following subset:

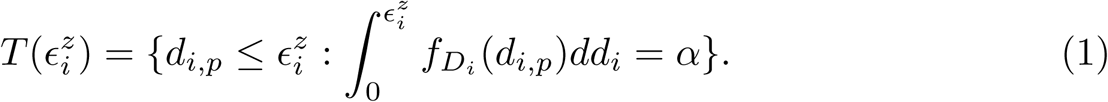

Therefore 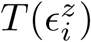 induces the subset 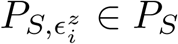 defined as:

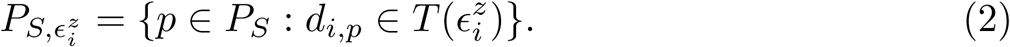

**5. Intersection of parameter sets.** Let denote 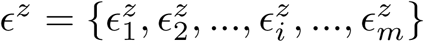 which is the set of thresholds corresponding to each observable at iteration *z*. We select the parameter samples that satisfy the conditions specified in the previous step simultaneously for all observables. This implies the definition of the following subset of *P*_*S*_:

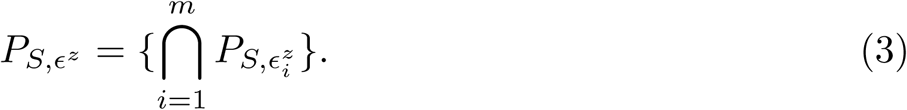

Thus, in the parameter space, the accepted samples belong to the approximate posterior distribution 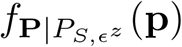. If the values of thresholds in *ϵ*^*z*^ satisfy the stopping criterion, the algorithm terminates and the output is 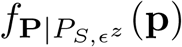. Otherwise go to step 6.

**6. Update the proposal distribution.** From 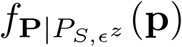 we select the mode vector 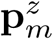. The proposal distribution of the subsequent iteration is defined as:

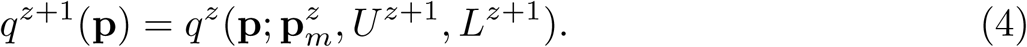

Note that the shape of the proposal distribution does not change over the different iterations, while the mean value is updated according to the mode 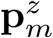 and the upper and lower boundaries, respectively *U*^*z*+1^ and *L*^*z*+1^, are shrinked through the following formulas:

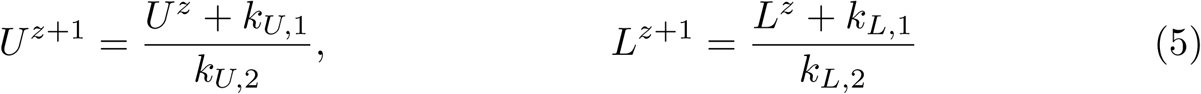

where *k*_*U,*1_, *k*_*U,*2_, *k*_*L,*1_ and *k*_*L,*2_ are constants set by the user. They regulate the shrinkage rate of the proposal distribution for all iterations performed. Once the new proposal distribution is defined, restart from step 1.

### Tuning parameters

In this section, we discuss the setting of the tuning parameters of the proposed algorithm. First of all, it is necessary to choose a sampling technique for the parameter space. The objective is to generate a fixed number *N*_*S*_ of samples that optimally cover the entire parameter space defined by the proposal distribution. To this purpose, we use Latin Hypercube Sampling (LHS) because it divides the multidimensional parameter space in 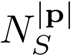 regions and guarantees that each region is represented by a single sample [26, 27]. Since several studies indicate that log-transforming the parameters usually yields a better performance, we use as sampling schema the Logarithmic Latin Hypercube Sampling (LoLHS) [28]. As for the number of samples *N*_*S*_, it is fixed in advance taking into account the dimension of the parameter space and the number of observables of the model.

Then, at each iteration the choice of the threshold values strictly depends on two constraints. First of all, in order to approximate the posterior distribution, tolerances have to be set so that 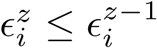. According to [22, 29], to generate a reliable non-parametric estimation of the conditional density, at least 1000 samples of 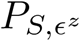 are necessary. Thus tolerances 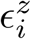 are chosen in order to ensure that the cardinality of 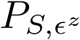 is at least 1000.

Moreover, different combinations of tolerances may lead to the fulfillment of this condition. Therefore, as a guideline, threshold values can be chosen so that the number of accepted samples for each distance function 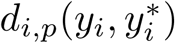 is similar as much as possible for all output variables. As the number of iteration increases, thresholds progressively shift toward zero and, as in the standard ABC-SMC method [30], this guarantees that the approximate posterior distribution 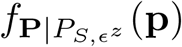 evolves toward the desired posterior distribution 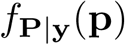.

The constraints explained above regarding the choice of the thresholds are also influenced by *N*_*S*_. For a given set of thresholds *ϵ*^*z*^, the higher the value of *N*_*S*_ the higher is the cardinality of 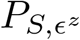. Thus, increasing *N*_*S*_, it is more likely to reach lower threshold values that satisfy 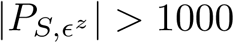. However, *N*_*S*_ has also great impact on the computational cost of CRC and, for these reasons, its choice is a trade-off between the accuracy of the posterior estimation and the efficiency of CRC.

Another tuning parameter of CRC is the definition of the distance function that is used to measure the level of agreement between simulations and experimental data.

Here, we propose two *l*1-*norm* based distance functions because they are robust to measurement noise [31].

The first one, called Absolute Distance Function (ADF), is the sum, over the whole time points set, of the absolute distance between simulated and real data. Eq (6) formalizes it:

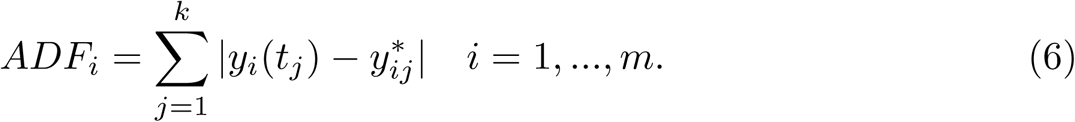

As an alternative, Eq (7) normalizes the absolute error between simulated and real data with the corresponding point in the dataset. Moreover, the summation is divided by the number of available time points, thus Eq (7) represents the mean percentage error on each time point.

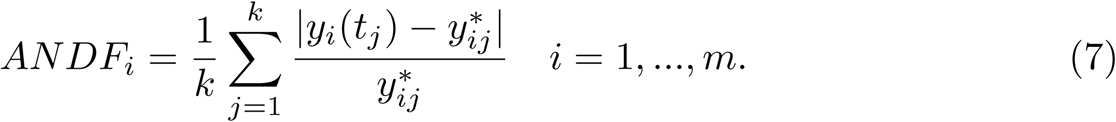

The criterion that guides the user in the choice of the best distance function regards the range of variation of the available data. For a given absolute error (Eq (6)), the corresponding percentage error (Eq (7)) produces a bias. Thus, Eq (7) is preferred when the model and data are normalized.

Finally, as explained above, parameters *k*_*U,*1_, *k*_*U,*2_, *k*_*L,*1_ and *k*_*L,*2_ in Eq (5) determine the contraction rate of the proposal distribution at each iteration. They should be chosen so that the proposal distribution is gradually shrinked iteration by iteration.

In CRC the number of necessary iterations is not known a priori because the algorithm terminates when the stopping criterion is satisfied.

### Conditional Robustness Analysis

The goal of the second phase of CRC is to perform a conditional robustness analysis (CRA) in order to identify which parameters mostly influence the output variables behavior [22]. To this purpose, we apply the conditional robustness algorithm in [22]. Starting from 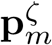 i.e., the mode in the parameter space obtained in the last CRC iteration, we sample the parameter space using Linear LHS and choosing as evaluation functions the distance functions 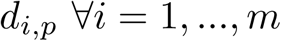 previously defined during the calibration process. For each distance function, we define two thresholds 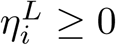 and 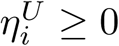 which are the quartiles of level *β* and *λ* respectively, obtaining the following subsets:

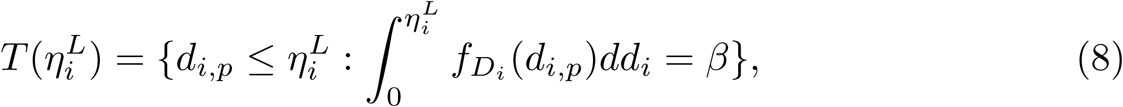

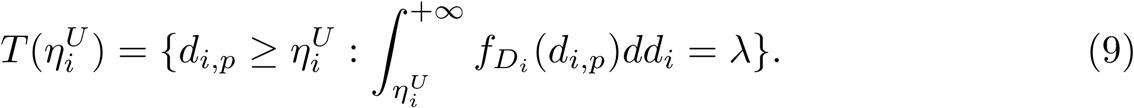

Thus, 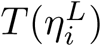 and 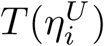 contain values of the distance functions 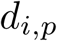 that are respectively smaller and larger than the defined thresholds 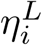 and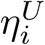. Therefore, 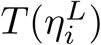 and 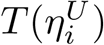 induce two subsets in the parameter space *P*_*S*_:

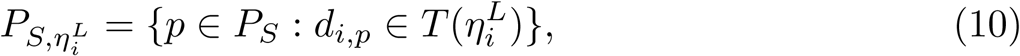

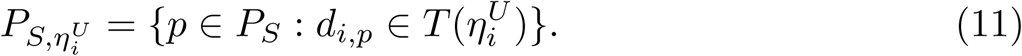

The resulting conditioning sets are:

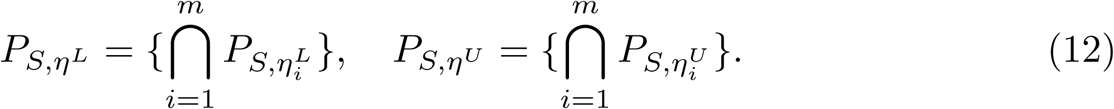

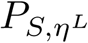 and 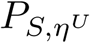 define two regions in the parameter space, whose samples belong to the distributions 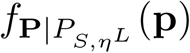 and 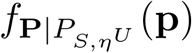.

The two conditional densities are employed in the calculation of the Moment Independent Robustness Indicator (MIRI) [22, 29] according to the following formula:

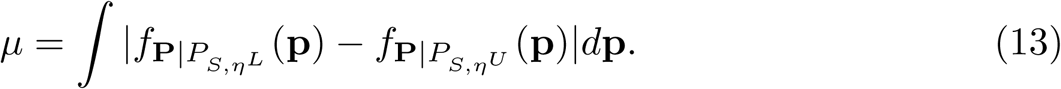

Vector *µ* contains MIRI values of all the components of parameter vector **p**. Due to its definition, the MIRI value of each parameter is included in the interval [0, 2]. MIRIs measure the level of intersection between the two pdfs included in the calculus: the higher the resulting value and the more well separeted are the conditional densities. Parameters with higher values of the MIRI have a major impact on output variables because there is a larger shift between the two conditional densities used for MIRI calculation. This means that different ranges of values for parameters with an high MIRI value lead the model observables to completely different behaviors. On the other hand, parameters with a low MIRI value do not affect the observables because their conditional densities overlap.

## Results

We test our novel proposed algorithm in three different models: Lotka-Volterra model (M1), EpoR system (M2) and signaling pathway of p38MAPK in multiple myeloma (MM) (M3). Table 1 synthesizes the models features. M1 is characterized by an oscillatory behavior of both output variables, M2 is used in synthetic biology and contains initial conditions and scale factors to estimate while M3 is a high-dimensional model based on experimental proteomics data.

**Table 1.** Features of the models.

| Model | Total parameters | Unknown parameters | States | Outputs | Data points |
| --- | --- | --- | --- | --- | --- |
| M1 | 4 | 2 | 2 | 2 | 8 |
| M2 | 11 | 7 | 5 | 2 | 22 |
| M3 | 93 | 53 | 40 | 16 | 96 |

### Lotka-Volterra model (M1)

#### Model description

The first model is the classical Lotka-Volterra model, which describes the interaction between the prey species *x*_1_ and the predator species *x*_2_ through the parameters *a* and *b*:

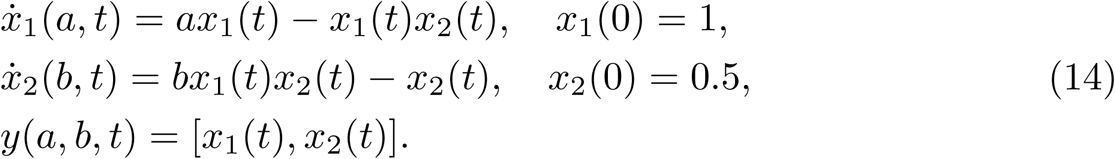

The model parameter vector to estimate is **p** = [*a, b*], **p** ∈ ℝ^2^. Both nominal values of parameters are set to 1. The observables are both variables *x*_1_ and *x*_2_.

To calibrate the model, we generate an *in silico* noisy dataset in the same way described by [30], i.e. sampling eight values of the output variables at the same specified time points and adding Gaussian noise 𝒩(0, (0.5^2^)) (Table S1 in S1 File).

#### CRC results

The tuning parameters of CRC are set as follows:

- the number of fixed samples in the parameter space is set to *N*_*S*_ = 10^4^ for each iteration;
- Eq (6) ∀*i* = 1, *…*2 is chosen as distance function between experimental and simulated data;
- the prior distributions for *a* and *b* are taken to be log-uniform: *a, b ∼ log − U* (0.1, 10);
- *k*_*U,*1_ = *k*_*L,*1_ = 1 and *k*_*U,*2_ = *k*_*L,*2_ = 2 in all iterations.

According to Eq (6), the distance between the noisy dataset and the nominal solution is 3.09 for species *x*_1_ and 2.6 for species *x*_2_. Thus, the objective of the calibration is to obtain realizations of 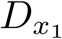 and 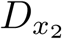 that are close, respectively, to 3.09 and to 2.6. For this reason, the algorithm terminates when the two corresponding thresholds become sufficiently close to these two values. In order to do that, we perform six iterations of CRC. In the sixth iteration 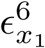 and 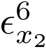 are very close to the reference values presented above (3.09 and 2.6). This demonstrates that CRC reaches the desired level of agreement between simulated and experimental data. Table 2 sums up the tuning parameters and the results of CRC along the six iterations.

**Table 2.** Tuning parameters and results of CRC in model M1.

| Iteration (z) | $L^z$ | $U^z$ | $\mathbf{p}_m^z$ | $\epsilon_{x_1}^z$ | $\epsilon_{x_2}^z$ | $MSE_{x_1}$ | $MSE_{x_2}$ |
| --- | --- | --- | --- | --- | --- | --- | --- |
| 1 | 0.1 | 10 | [0.37, 0.96] | 7.3 | 6 | 0.71 | 0.54 |
| 2 | 0.55 | 5.5 | [0.81, 1.26] | 5.8 | 4 | 0.38 | 0.2 |
| 3 | 0.775 | 3.25 | [0.84, 1.11] | 5 | 3.4 | 0.35 | 0.19 |
| 4 | 0.8875 | 2.125 | [0.98, 1.07] | 4 | 3 | 0.22 | 0.16 |
| 5 | 0.9437 | 1.5625 | [1.01, 1.07] | 3.4 | 2.7 | 0.2 | 0.14 |
| 6 | 0.9718 | 1.2813 | [1.06, 1.06] | 3.1 | 2.7 | 0.19 | 0.13 |

The second and third column show, respectively, the lower and upper boundaries of $q^z(\mathbf{p})$ . The fourth column reports the mode vector $\mathbf{p}_m^z$ computed at the end of each iteration and used to center the proposal distribution of the subsequent iteration. In the fifth and sixth columns the threshold schedule is reported and in the last two columns the resulting MSE for each observable is shown.

Fig 2A and Fig 2B display the time behavior of output variables *x*_1_ and *x*_2_ when parameters belong to the final subset 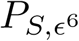. The model simulations are shown together with the noisy dataset, proving both the validity and robustness of the solution. Fig 2C shows how the subset of accepted particles 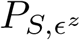 changes during the execution of CRC.

**Fig 2.**
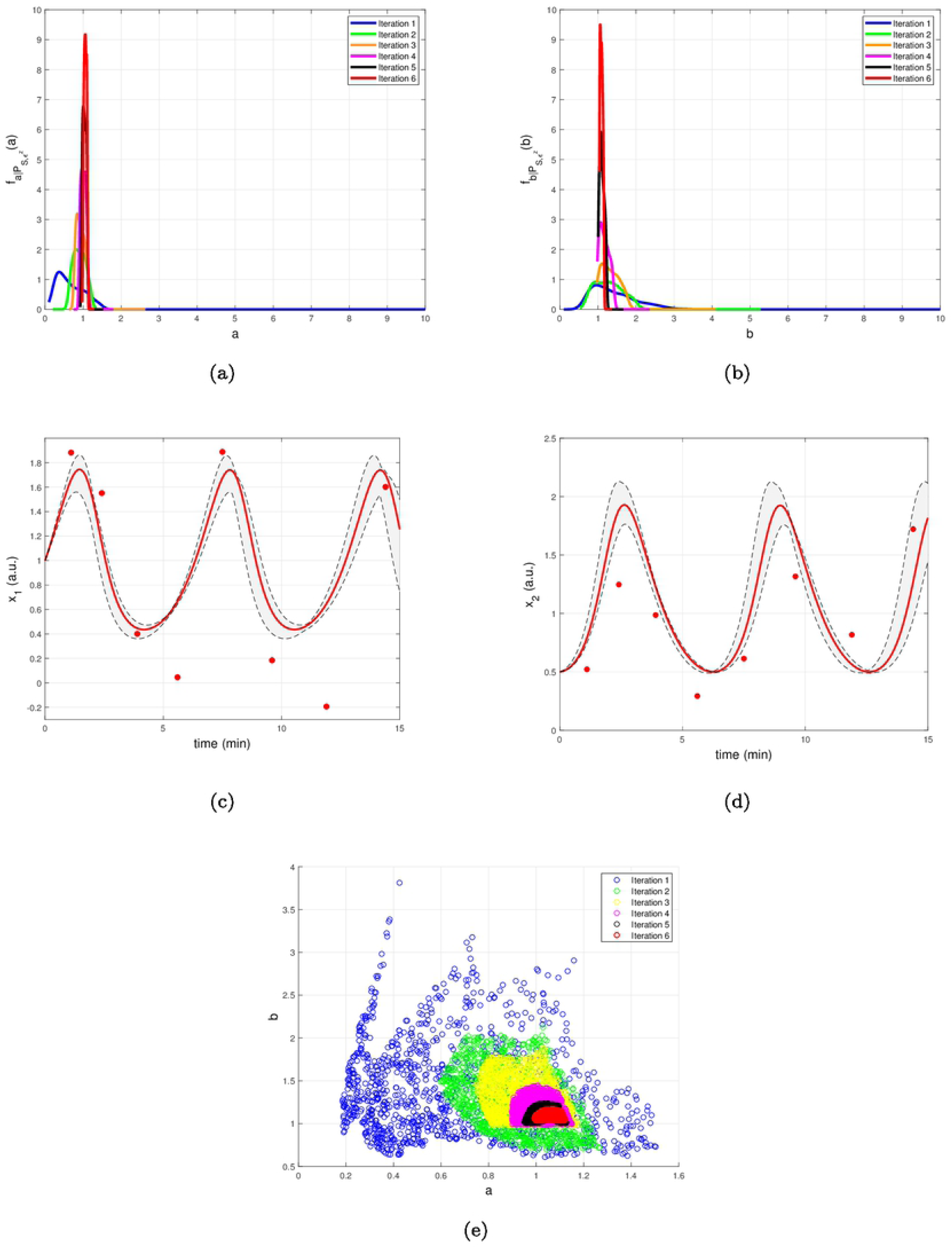
Results of CRC for model M1. (A), (B) The red line is the time behavior of output variables when the parameter vector is equal to **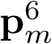** (see Table 2); red dots are the experimental data; the gray area represents the uncertainty in the temporal behavior of observables when parameters vary between the 2.5th and 97.5th percentile of their corresponding conditional pdfs. (C) Scatter plot of the sample distribution of 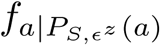 (x-axis) versus 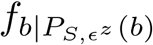 (y-axis) in all iterations.

CRC is fast in model calibration since the time simulation decreases from 288 seconds (s) for the first iteration until 202 s for the last one. After model calibration, the CRA is applied in order to compute MIRIs for the parameters. We perturb the mode vector 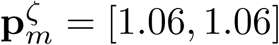 generating 10^4^ samples of the parameter space. The lower and upper boundaries of the sampling are fixed equal to 0.1 and 10 and the probabilities *β* and *λ* are both fixed to 0.1, following the guidelines in [22]. For both parameters, we obtain that MIRIs have values around 1, meaning that *a* and *b* have approximately the same influence on the two output variables (Fig S3 in S1 File).

#### PL results

First of all, we estimate parameter values using three different optimization algorithms, available in the software D2D [13]: *lsqnonlin*, genetic algorithms (GA) and simulated annealing (SA). In this example, both the default *lsqnonlin* and GA correctly fit the model yielding the same results, while SA totally fails in parameter estimation. The default method *lsqnonlin* estimates parameter *a* equal to 1.07 and parameter *b* equal to 1.05. Using these parameter values, the MSE is equal, respectively, to 0.19 for *x*_1_ and to 0.13 for *x*_2_. In order to evaluate the identifiability of model parameters and to assess their confidence intervals, we also calculate the PL. All tuning parameters of the algorithm are left to their default values. PL estimates as identifiable both model parameters (Fig S5 in S1 File). PL employs few seconds for parameter estimation and less than a minute for identifiability analysis of both parameters.

#### ABC-SMC results

The application of the standard ABC-SMC method to the M1 model and the results obtained are comprehensively explained in [30]. In five steps the procedure converges to the considered threshold. In the *5-th* iteration, parameter *a* has a median of 1.05 and a 95% interquartile range of [1, 1.12] while parameter *b* has a median of 1 and a 95% interquartile range of [0.87, 1.11] [30]. When parameters are equal to their median values, 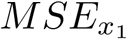 is 0.28 and 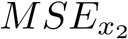 is 0.19.

#### DRAM results

We apply DRAM to the M1 model varying the initial parameter values in the interval [0, 10], the initial error variance in the interval [0.01, 1] and the corresponding prior weight in the interval [1, 5]. Moreover, in order to perform a reliable comparison with CRC, we set the number of simulations equal to 10^4^ and the boundaries of the parameters between [0, 10]. In all cases, the chains are stable and close to the nominal parameter values after one run. For the sake of brevity, we show only the results for initial error variance set to 0.1, prior weight set to 3 and initial parameter values equal to 0, 5 and 10. The time employed to apply the algorithm on the M1 model is about 3 minutes. Detailed results of DRAM are in the S1 File.

### EpoR System (M2)

#### Model description

The ODE model presented in this section is taken from the Erythropoietin Receptor (EpoR) [32]. The model represents the catalysation of a substrate S by an enzyme E that is activated via two steps by an external ligand L [33]. This reaction cascade produces a product P whose dynamical behavior is the purpose of the model prediction. Generally the concentration over the time of the product P cannot be measured directly. Let denote with **p** = [*k*_1_, *k*_2_, *k*_3_, *init*_*E*_, *init*_*S*_, *scale*_*E*_, *scale*_*S*_], **p** ∈ ℝ^7^, the set of parameter to estimate. The nominal values of model parameters are **p** = [0.1, 0.1, 0.1, 10, 5, 4, 2]. The equations of the model, the corresponding initial conditions and the observables together with the dataset used for model calibration are reported in S1 File.

#### CRC results

The prior distributions for all the model parameters are supposed log-uniform with the lower and upper boundaries set equal to *L*^1^ = 0.01 and *U* ^1^ = 100. The number of fixed samples in the parameter space is *N*_*S*_ = 10^5^. We choose Eq (6) as distance function to evaluate the error between nominal and noisy data for the outputs of the model. According to the selected distance function, the errors between the nominal data points and the experimental ones are equal to 12.78 for *y*_1_ and 5.6 for *y*_2_. They represent the target thresholds to reach at the end of the last iteration in order to assert the success of CRC. To this purpose, we perform nine iterations of CRC in order to make the two thresholds close enough to their corresponding target values. Table 3 shows the boundaries of the proposal distribution in each iteration. These values are obtained by setting *k*_*U,*1_ = *k*_*L,*1_ = 0, *k*_*U,*2_ = 2 and *k*_*L,*2_ = 0.5 for the first seven iterations and *k*_*U,*1_ = *k*_*L,*1_ = 1 and *k*_*U,*2_ = *k*_*L,*2_ = 2 for the eighth and ninth iterations. Table 3 also shows the obtained thresholds for each performed iteration of CRC.

**Table 3.** CRC parameters for model M2.

| Iteration ($z$) | $L^z$ | $U^z$ | $\epsilon_1^z$ | $\epsilon_2^z$ |
| --- | --- | --- | --- | --- |
| 1 | 0.01 | 100 | 51 | 31 |
| 2 | 0.02 | 50 | 50 | 31 |
| 3 | 0.04 | 25 | 46 | 30.9 |
| 4 | 0.08 | 12.5 | 40 | 30.3 |
| 5 | 0.16 | 6.25 | 33.5 | 29.1 |
| 6 | 0.32 | 3.125 | 25 | 27.5 |
| 7 | 0.64 | 1.5625 | 15.5 | 22 |
| 8 | 0.82 | 1.2813 | 13.4 | 10 |
| 9 | 0.91 | 1.1406 | 13 | 5.75 |

The first column reports the iteration number $z$ , the second and the third ones the boundaries of the proposal distribution $q^z(\mathbf{p})$ in each iteration and the fourth and the fifth ones the values of the two thresholds $\epsilon_1^z$ and $\epsilon_2^z$ .

In the ninth iteration, 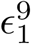 and 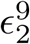 are very similar to the target values presented above. This proves that CRC estimates a parameter vector that guarantees the desired level of agreement between simulated and experimental data. The mode vector in output from CRC is 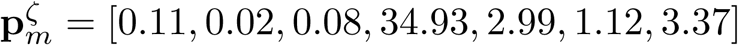. The model simulation using as parameter vector the mode 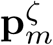 has an MSE of 18.24 and 0.32 for *y*_1_ and *y*_2_ respectively. Fig 3A and Fig 3B show the time behavior of both output variables when the parameter vector is set equal to the mode 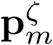.

**Fig 3.**
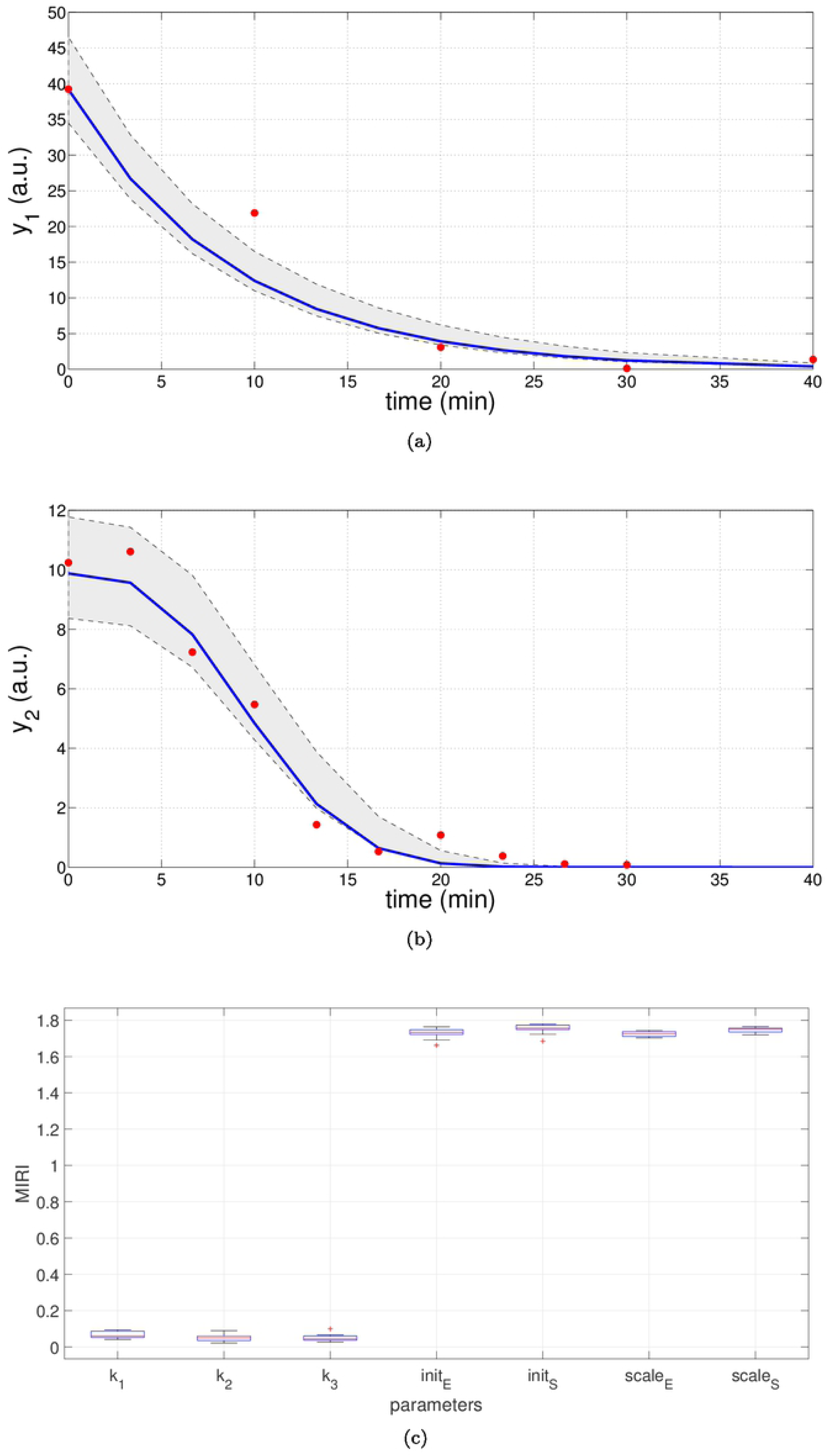
Results of CRC for model M2. (A), (B) Time behavior of output variables. Red dots are the noisy experimental data [33]); blue lines are the simulations of the observables of the model when the parameter vector is set equal to the mode 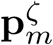; gray regions are the confidence bands when parameter values are chosen between the *2.5-th* and *97.5-th* percentile of their corresponding conditional pdfs 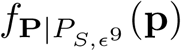. (C) Boxplots of the MIRIs in output from the CRA for the seven model parameters and all the ten independent realizations.

Moreover, regions in gray are the confidence bands of observables when parameter values are chosen between the *2.5-th* and *97.5-th* percentile of their corresponding conditional pdfs 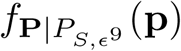. In S1 File additional details of 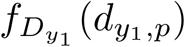 and 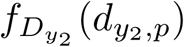 and the estimated conditional pdfs of parameters are reported. CRC is quite fast since it employs about 8 minutes (527 s) to complete one iteration.

Once the model has been correctly calibrated, we perform a robustness analysis in order to find those parameters that most affect the behavior of the output variables. We perturb the mode vector 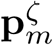 with Linear LHS using 10^5^ samples. The lower and upper boundaries of the sampling are fixed equal to 0.01 and 100 respectively. Using the guidelines reported in [22] we fix the level of probabilities *β* and *λ* to 0.1. In Fig 3C the resulting MIRIs are shown. MIRIs corresponding to initial conditions parameters and scale factors are close to their maximum value and are much higher than those of the kinetic ones. This means that initial conditions and scale factors have major impact on observables compared to the kinetic parameters. We repeat the entire procedure ten times, obtaining ten independent realizations in order to ensure the invariance of results.

#### PL results

The calculation of PL for M2 is presented in [33]. PL takes less than one minute per parameter on a 1.8 GHz dual core machine. The parameter vector estimated through the PL is 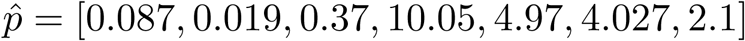. The MSE obtained through the PL approach is 10.06 and 0.3 for *y*_1_ and *y*_2_ respectively. As regards the identifiability analysis, according to the PL approach, parameter *k*_2_ is classified as structurally non-identifiable, parameter *k*_3_ is practically non-identifiable and the others are assessed to be identifiable. More details of the PL results are provided in S1 File.

#### ABC-SMC results

ABC-SMC input parameters are set in order to resemble those of CRC. The distance function is defined as:

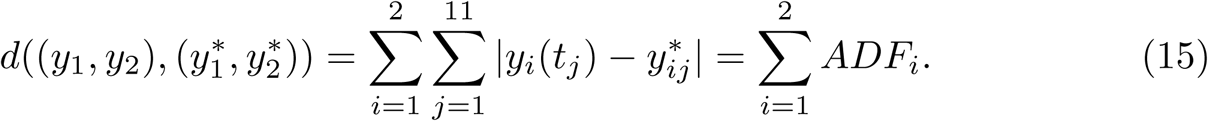

Under the hypothesis of parameters having a prior uniform distribution in [0, 100], we try to perform nine iterations of ABC-SMC. Thus, we set *f***_P_**(**p**) = *U* (0, 100) and *z* = 1, *…,* 9. The thresholds for all the iterations are chosen as the sum, over *y*_1_ and *y*_2_, of the two corresponding thresholds obtained from the application of CRC (Table 3). At each iteration, we select 1000 particles of the parameter space under the desired threshold. The algorithm could not come to an end in a reasonable time and it finds parameter samples only until the *7-th* iteration. In order to quantify the precision of the ABC-SMC approach, we perform simulations of the model, setting the parameter vector equal to the median of the ABC-SMC results. The MSE using these parameter values is 103 and 20 for *y*_1_ and *y*_2_ respectively. Further results of ABC-SMC are shown in S1 File.

#### DRAM results

As in model M1, we run DRAM varying the initial error variance between [0.01,1] and the corresponding prior weight between [1, 5]. Initial conditions were chosen close to nominal parameter values, i.e. [0.5, 0.5, 0.5, 8, 7, 5, 3]. As in CRC, the number of simulations is set to 10^5^ and the parameter boundaries are [0, 100]. In all cases, the chains do not converge, even when performing multiple iterative runs of DRAM. Here, we report the results obtained with a prior weight set to 1 and an initial error variance that varies between 0.01 and 1. DRAM employs about 15 minutes to complete one run. In S1 File figures of DRAM results are provided.

### Multiple myeloma model (M3)

#### Model description

M3 is the ODE model proposed in [34]. The mathematical model is defined to help study the roles that various p38 MAPK isoforms play in MM. It has 40 ODEs, built using only the law of mass action, and 53 kinetic parameters (S1 File). According to [34], data associated to the model are the output of a Reverse Phase Protein Array (RPPA) experiment [2], where MM cell lines were analyzed to detect the activity of proteins active in various p38 MAPK signaling pathways. RPPA was performed on the following cell lines: four different RPMI 8226 MM cell sublines with stable silenced expression of *p*38*α, p*38*β, p*38*γ* and *p*38*δ*, respectively, as well as an RPMI 8226 MM stable cell subline transfected with empty vector as the negative control. Cells were treated with arsenic trioxide (ATO), bortezomib (BZM) or their combination. RPPA analyzed a total of 80 samples and measured 153 proteins at six different time points. The proteins whose phosporylation level was included both in the pathway and in the experiment are 16. All RPPA data are normalized based on the initial concentration value. For this reason, initial conditions of proteins in the ODE model are all set to 1, i.e. **x**(0)=**1**. Parameters to estimate are only the kinetic ones, i.e. **p** ∈ ℝ^53^. In [34], available data for model calibration belongs to the *p*38*δ* knockdown cell line treated with BZM. The corresponding RPPA dataset is presented in Table S13 of S1 File.

#### CRC results

For model M3, we set tuning parameters of CRC in the following way:

- the number of parameter samples *N*_*S*_ is equal to 10^6^;
- for each output variable, Eq (7) is chosen as distance function;
- the prior of each kinetic parameter is supposed to be *log - U* (0.1, 10);
- *k*_*U,*1_ = *k*_*L,*1_ = 1 and *k*_*U,*2_ = *k*_*L,*2_ = 2 in all iterations.

Six iterations of CRC are performed, using the threshold schedule reported in Table 4. At the end of the process, the maximum error between simulated and experimental data is only of the 16%.

**Table 4.** Threshold schedule of distance functions for model M3.

| | Iteration ($z$) | | | | | |
| --- | --- | --- | --- | --- | --- | --- |
|  | 1 | 2 | 3 | 4 | 5 | 6 |
| *RasGTP* | 0.7 | 0.4 | 0.3 | 0.2 | 0.1 | 0.08 |
| *pPI3K* | 0.7 | 0.4 | 0.3 | 0.2 | 0.1 | 0.06 |
| *pp38* | 0.7 | 0.4 | 0.3 | 0.15 | 0.13 | 0.09 |
| *pPDK1* | 0.8 | 0.5 | 0.3 | 0.2 | 0.1 | 0.06 |
| *pAKT* | 0.3 | 0.2 | 0.15 | 0.12 | 0.095 | 0.088 |
| *pmTOR* | 0.5 | 0.3 | 0.2 | 0.13 | 0.1 | 0.088 |
| *pRaf1* | 0.7 | 0.4 | 0.2 | 0.15 | 0.1 | 0.08 |
| *pMEK12* | 0.7 | 0.4 | 0.3 | 0.2 | 0.1 | 0.08 |
| *pERK12* | 0.6 | 0.4 | 0.2 | 0.1 | 0.05 | 0.04 |
| *pP70S6k* | 0.5 | 0.3 | 0.2 | 0.15 | 0.1 | 0.08 |
| *pcJUN* | 0.6 | 0.3 | 0.2 | 0.15 | 0.12 | 0.12 |
| *BCLXL* | 1 | 0.7 | 0.5 | 0.3 | 0.2 | 0.15 |
| *BAX* | 0.8 | 0.6 | 0.5 | 0.35 | 0.2 | 0.15 |
| *pNFkB* | 0.6 | 0.4 | 0.25 | 0.2 | 0.18 | 0.16 |
| *cPARP* | 0.7 | 0.4 | 0.3 | 0.2 | 0.15 | 0.12 |

The distance functions are $d_{i,p}(y_i, y_i^*)$ , $\forall i = 1, \dots, 16$ .

Fig 4A shows the time behavior of output variables at the end of the calibration, along with experimental data points. The figure proves that the algorithm is successful in finding a robust solution (S1 File). As regards the application of CRA, we perturb the parameter space in the interval having as lower and upper boundaries 0.01 and 100 respectively and centered around the final mode vector 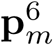, generating 10^6^ parameter samples. In Fig 4B, MIRIs are presented for each parameter. As expected, not all parameters have a strong impact on the observables since the number of output variables is small compared to all kinetic parameters. For instance, parameters *k*_*Raf*1___*pPI*3*K*_, *k*_*ERK*12_*pp*38_, *k*_*BAX_pp*38_, *k*_*IKK_pAKT*_, *k*_*PARP_BAX*_, *k*_*cPARP_BCLXL*_ have all a MIRI value under 0.2. On the other hand, about a fifth of the total number of parameters has MIRI values above 1.6 (e.g. parameters *k*_*GFR*_, *k*_*pGFR*_, *k*_*Shc_pGFR*_, *k*_*pPI*3*K*_, *k*_*pAKT*_, *k*_*pRaf*1_*pAkt*_, *k*_*pMEK*12_, *k*_*ERK*12_*MEK*12_, *k*_*pERK*12_, *k*_*pP*70*S*6*K*_, *k*_*pJNK*_), meaning that they have high influence on the outputs. We repeat the entire procedure ten times, obtaining ten independent realizations in order to ensure the invariance of results.

**Fig 4.**
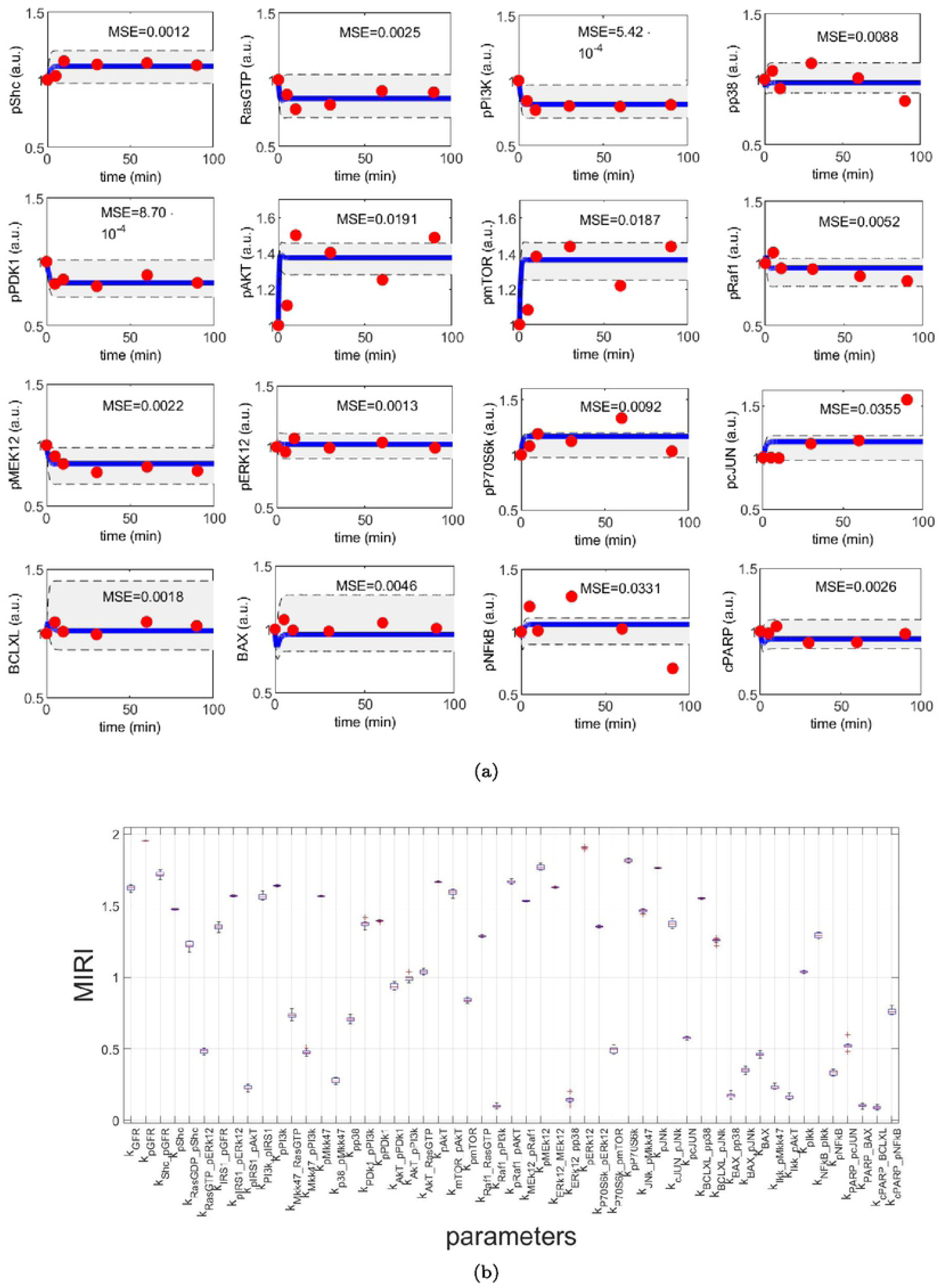
CRC results for model M3. (A) The blue line is the time behavior of observables when the parameter vector is equal to **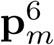** (see Table S25 in S1 File); red dots are the RPPA data [34]); the gray area reproduces the variation of the temporal behavior when parameters vary between the *2.5-th* and *97.5-th* percentile of their corresponding conditional pdfs 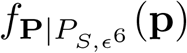. On the top of each plot, the MSE between the associated model simulation and the data is reported. (B) Boxplot of MIRIs for the estimated parameter values, along the ten independent realizations performed.

#### PL results

First of all, through the D2D software, we calibrate the model using *lsqnonlin* as optimization algorithm. To avoid local optima, we execute a sequence of *n* = 100 fits using Linear LHS. Moreover, we also estimate model parameters using GA and SA. While GA is not able to correctly reproduce experimental data, SA successfully estimates the time behavior of output variables. To compute confidence intervals, the maximum number of sampling steps in both the increasing and decreasing direction of each parameter is set to 200 while all the other tuning parameters of the method are set to their default values. Results of parameter estimation and identifiability analysis are shown in S1 File. The algorithm assesses that all parameters are identifiable except for the following parameters: *k*_*P*70*S*6*K*___*pϵRK*12_, *k*_*cPARP*___*BCLXL*_, *k*_*pJNK*_ that are practically non-identifiable and *k*_*pIRS*1___*pAKT*_ is structurally non-identifiable. Nevertheless, the results obtained are not so reliable since some parameters have a confidence interval of only a single value (e.g. *k*_*IRS*1___*pGFR*_) while others have an estimated value outside the corresponding confidence region (e.g. *k*_*P DK*1___*pPI*3*K*_). The PL algorithm employs less than one minute for parameter estimation and about 40 minutes for identifiability analysis of all parameters.

#### ABC-SMC results

Using the ABC-SysBio software, we fix ABC-SMC parameters as follows:

- the distance function is defined as:

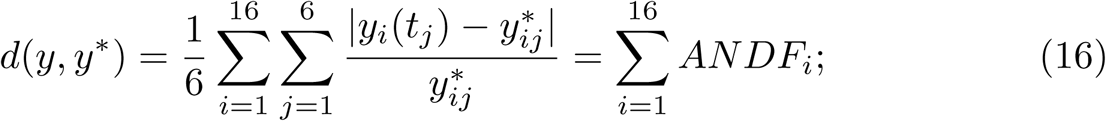
- the number of iterations is set to 6;
- the threshold fixed in each iteration is equal to the sum of all thresholds fixed in CRC in the same iteration (see Table S29 of S1 File);
- the number of accepted particles at each iteration is set to 1000;
- all parameters to estimate are supposed to have a uniform prior distribution: U(0.1,10);
- all the other parameters are left to their default values.

The application of ABC-SMC to the M3 model was very time consuming and, after 10 days, it had not converged yet. Thus, results are available only until the *4-th* iteration (see S1 File).

#### DRAM results

As in the previous examples, we run DRAM setting the number of simulations equal to those of CRC (10^6^). The initial error variance is set to 0.1 and the corresponding prior weight to 10. Initial values of parameters are all equal to 1 and the parameter boundaries are [0, 10]. DRAM employs about 24 hours to complete one run and all chains of the parameters do not converge. In S1 File figures of DRAM results are provided.

## Discussion

Here we present a novel Bayesian approach for parameter estimation of mathematical models that is used to fit *omics* data in Systems Biology applications. The availability of high-throughput data with the need to calibrate high dimensional models using computational feasible algorithms, makes CRC a useful and innovative procedure in the overview of the Bayesian parameter estimation and robustness analysis. Our algorithm modifies the standard ABC-SMC in order to increase the efficiency and the reliability of the estimated parameter vector. Moreover CRC presents many distinctive improvements as compared to other algorithms of the ABC-SMC family, such as ABC-PMC and Adaptive-ABC [21].

We validated this new methodology in three ODE models, each one with specific features, in order to demonstrate the flexibility and reliability of our approach. In addition, we compared CRC results with those obtained by methods representing the state of the art of this field, i.e. the standard ABC-SMC, PL and DRAM.

First of all, we tested all the calibration procedures in the Lotka-Volterra model. We showed that all algorithms performed well, but with some differences. CRC returns a reliable and robust solution. Compared with ABC-SMC, we performed one more iteration but the computational burden was almost irrelevant since each iteration took about 5 minutes to complete. CRC also finds a more precise solution and generates a remarkable minor number of particles for sampling the parameter space. PL succeeds in fitting the data when *lsqnonlin* and GA are used. Then, through confidence intervals, it classifies both parameters as identifiable, in accordance with MIRI values. DRAM, after the initial adaptation period, finds acceptable points and a good mixing of the chain, regardless of the choice for the tuning parameter values.

Next we compared the results of CRC in the model presented in [32, 33]. In this example, CRC finds an alternative solution of the parameter vector compared to that of PL. PL fits the data properly and fast and identifiability results are in accordance with MIRI values. ABC-SMC fails in the calibration procedure and it cannot go beyond the *7-th* iteration, proving that it cannot reach an error as low as the one of CRC. As regards DRAM, even though we tried multiple combinations of the freely tuning parameters, the chains of the parameters do not converge and thus the method is unable to provide reliable results.

Finally, the last model is a high-dimensional ODE model calibrated on real experimental data [34]. CRC was able to find a set of parameter vectors that fit well experimental data. In addition, robustness analysis highlights that about half of the parameters influences most output variables. PL is successful in model calibration but computation of confidence intervals gives confounding results which do not allow a reliable comparison with MIRI values. ABC-SMC fails in model calibration because it remains blocked in the *5-th* out of 6 iterations. Also DRAM does not find a reliable solution since all the parameter chains are not stable and, after 10^6^ simulations, they span the interval [0, 10] almost uniformly.

In summary, the main disadvantage of the standard ABC-SMC method is the time necessary to complete a simulation which increases with the model dimension. The PL method is fast in model calibration even for high dimensional models since it implements an optimization algorithm. However, the returned solution does not contain any information on the distribution of parameters since it represents a single point in the parameter space. Moreover, as shown in model M3, it may return improper results in the computation of parameter profiles. As regards DRAM, its results are highly affected by the initial values of the parameters, which must be set from the beginning. This point is crucial since in Systems Biology models most parameter values are unknown and cannot be measured experimentally.

CRC is able to identify a stable and precise solution in all test models, mainly because of some of distinctive features. One of its main innovations is the use of a fixed number of points for sampling the parameter space, which is initially chosen by the user and does not change throughout iterations. As a result, the model is always integrated *N*_*S*_ times in each iteration. Since most of the computational cost of an iteration of CRC is given by the integration of the model (step 2 of CRC), CRC guarantees a limited computational cost through the different iterations. On the other hand, in other ABC-SMC methods, the computational burden is substantial because the number of samples at each iteration is not known *a priori* but strictly depends on the threshold value. Since the threshold usually decreases at each step, the number of generated samples could increase together with the simulation time.

Moreover, another significant innovation introduced is the definition of an objective function for each output variable. This allows a model calibration that takes equally into account all experimental endpoints. On the other hand, the other techniques evaluate only a single and unique objective function, which includes information about all observables.

Finally, we also analyzed the robustness of model parameters in a new way, taking inspiration from the CRA presented in [22]. This algorithm is based on the concept of robustness proposed by [35], which defines it as the property of a system to maintain its status against internal and external perturbations. We employed CRA in order to quantify the robustness of the model observables against the simultaneous perturbation of the parameters.

Robustness analysis is useful for applications in cancer drug discovery aimed at finding which node of a network could be identified as novel potential drug target. Moreover, the concept of robustness is slightly different from that of identifiability introduced with the PL approach. A parameter that is declared identifiable should have an high MIRI value since it has great impact on the outputs behavior. On the other hand, if a parameter is non-identifiable it is impossible to understand its influence on the observables dynamical response, without performing our robustness analysis. While ABC-SMC evaluates parameter identifiability only through histograms of final parameter values and DRAM computes the parameter posterior distribution, CRC estimates conditional parameter densities and performs robustness analysis through the MIRI indicator that quantifies the influence of each parameter on the behavior of interest. Indeed, the higher the MIRI value the higher the impact of the parameter on the entire set of observables. All the innovations introduced with CRC are important for a successful calibration of high dimensional nonlinear models in Systems Biology applications based on *omics* data.

## Supporting information

### S1 File. Supplementary methods and results

Description of the PL, DRAM and ABC-SMC methods. For each model equations and data are shown together with additional tables and figures of the results.

